# Fast and efficient QTL mapper for thousands of molecular phenotypes

**DOI:** 10.1101/022301

**Authors:** Halit Ongen, Alfonso Buil, Andrew Anand Brown, Emmanouil T. Dermitzakis, Olivier Delaneau

**Affiliations:** Department of Genetic Medicine and Development, University of Geneva Medical School, Geneva, Switzerland; Swiss Institute of Bioinformatics, Geneva, 1211, Switzerland

## Abstract

**Motivation:** In order to discover quantitative trait loci (QTLs), multi-dimensional genomic data sets combining DNA-seq and ChiP-/RNA-seq require methods that rapidly correlate tens of thousands of molecular phenotypes with millions of genetic variants while appropriately controlling for multiple testing.

**Results:** We have developed FastQTL, a method that implements a popular cis-QTL mapping strategy in a user- and cluster-friendly tool. FastQTL also proposes an efficient permutation procedure to control for multiple testing. The outcome of permutations is modeled using beta distributions trained from a few permutations and from which adjusted p-values can be estimated at any level of significance with little computational cost. The Geuvadis & GTEx pilot data sets can be now easily analyzed an order of magnitude faster than previous approaches.

**Availability:** Source code, binaries and comprehensive documentation of FastQTL are freely available to download at http://fastqtl.sourceforge.net/.

## Introduction

Genome-wide association studies have shown that most common trait-associated variants fall into non-coding genomic regions and likely alter gene regulation (Maurano et al., 2012; Nica et al., 2010). This has motivated large scale studies to catalog candidate regulatory variants (Quantitative Trait Loci; QTL) associated with various molecular phenotypes (i.e. quantitative molecular traits with a genomic location) across various populations (Lappalainen et al., 2013), cell (Fairfax et al. 2012) and tissue types (GTEx Consortium., 2015; Ongen et al., 2014). Mapping QTLs in this context usually consists of finding statistically significant associations between phenotype quantifications and nearby genetic variants. This requires millions of association tests in order to scan all possible phenotype-variant pairs in cis (i.e. variants located within a specific window around a phenotype), resulting in millions of nominal p-values. Matrix eQTL (Shabalin., 2012) has recently emerged as a “gold standard” for this task (Lappalainen et al., 2013; GTEx Consortium., 2015) by taking advantage of efficient matrix operation implementations to perform the many association tests in acceptable running times. Due to the large number of tests performed per phenotype, multiple testing has to be accounted for to assess the significance of any discovered candidate QTL. A first naive solution to this problem is to correct the nominal p-values for the number of tested variants using the Bonferroni method. However, due to the specific and highly variable nature of each genomic region being tested in terms of allele frequency and linkage disequilibrium (LD), the Bonferroni method usually proves to be overly stringent and results in many false negatives. To overcome this issue, a commonly adopted approach (Montgomery et al., 2010) is to analyze thousands of permuted datasets for each phenotype in order to empirically characterize the null distribution of associations (i.e. the distribution of p-values expected under the null hypothesis of no associations). Then, we can easily assess how likely an observed association obtained in the nominal pass originates from the null, resulting in an adjusted p-value. In practice, performing permutations in this context requires fast methods able to absorb such substantial computational loads in reasonable running times. Even though Matrix eQTL has been used so far in multiple large scale studies (Lappalainen et al., 2013; GTEx Consortium., 2015), it still suffers from a main drawback which makes its practical application relatively laborious and time consuming: there is no efficient built-in permutation scheme forcing users to develop their own and therefore to use non-optimal multiple-testing correction methods. So far, a commonly employed permutation strategy relies on performing a fixed number of permutations per phenotype (1,000 to 10,000) to control the running times at the cost of accurately assessing the statistical significance of the most strongly associated QTLs. Here we present FastQTL, a user- and cluster-friendly QTL mapper, which improves upon Matrix eQTL by implementing a fast and efficient permutation scheme in which the null distribution of associations for a phenotype is modeled using a beta distribution. This allows us to approximate the tail of the null distribution relatively well using only few permutations, and then to accurately estimate adjusted p-values at any significance level in short running times.

## Methods

### 1. Overview

FastQTL performs linear regressions between genotypes and molecular phenotypes with or without covariates in order to find the best nominal association for each phenotype (Methods Section 2). Then, it can correct for the multiple correlated variants tested via three different permutation schemes: (1) a direct permutation scheme that relies on a fixed number of permutations (Methods Section 3), (2) an adaptive permutation scheme which maintains a reasonable computational load by tailoring the number of permutations to the significance of the association (Methods Section 4) and (3) a beta approximation which models the permutation outcome via a beta distribution (Methods Section 5). For (1) and (2), an adjusted p-value per phenotype is calculated as the proportion of null associations found to be more significant than the nominal one. For (3), we model this null distribution of most significant p-values for a phenotype with a beta distribution, learning the parameters from a few permutations (typically 100 to 1,000) by maximum likelihood estimation (MLE). As a result, we obtain a reasonably good approximation of the tail of null distribution to estimate small adjusted p-values at any significance level (i.e. without lower bound). In a final stage, a false discovery rate (FDR) procedure as implemented in the R/qvalue (Storey and Tibshirani., 2003) package is used on the set of adjusted p-values obtained either from (1), (2) or (3) to extract all significant phenotype-variant pairs at a given FDR, usually chosen to be 5 or 10% (Methods Section 6). All this, plus other optional functionalities, have been implemented in the FastQTL software package (Methods Section 7).

### 2. Finding a candidate QTL per phenotype

For simplicity, we will focus on a single molecular phenotype **P** quantified in a set of **N** samples. Let **G** be the set of genotype dosages at **L** variant sites located within a cis-window of +/- **W** Mb of the genomic location of **P**. To discover the best candidate QTL for **P**, FastQTL measures Pearson product-moment correlation coefficients between **P** and all **L** variants in **G**, stores the most strongly correlated variant **q □ G** as candidate QTL, and assesses its significance by calculating a nominal p-value **p_n_** with standard significance tests for Pearson correlation. Note that this is equivalent to testing for **β ≠0** in a linear model **P = βg + E** with **β** estimated by least squares fitting. Of note, this method is also used by Matrix eQTL to speed up linear regression (Methods 3.1 & 3.2 of Shabalin., 2012). Then, two multiple-testing levels are accounted for to determine the whole-genome significance of this nominal p-value and thereby to consider the corresponding variant **q** as a QTL: multiple genetic variants are tested per phenotype and multiple phenotypes are tested genome-wide. Following common usage, FastQTL uses permutations to correct for the former (Methods Sections 3-5) and false discovery rate estimation to control for the latter (Methods Section 6).

### 3. Direct permutation scheme

The choice of a proper global significance threshold for nominal p-values is very difficult due to the fact that we test multiple variants per phenotype, correlated because of linkage disequilibrium (LD), across a wide allele frequency spectrum, whilst all this varies from one phenotype to another. To account for this, significance of a candidate QTL is assessed via permutations. Specifically, we repeat the cis-window scan procedure for **R** random permutations of **P**, leaving the genotype data **G** unchanged to preserve the correlation between variants. Each time, we store the strongest correlation; the goal is to produce a sample from the distribution of the strongest correlation under the complete null hypothesis of no genetic associations. Then, the observed correlation is compared to this empirical null distribution to obtain an adjusted p-value characterizing the significance of the candidate QTL. When very few null correlations are found to be stronger than the observed one, it means that reaching this correlation level by chance is very unlikely and therefore that the QTL candidate is likely to be true. More formally, if **r** correlations in the null distribution are found to be stronger than the observed, significance of the QTL candidate is assessed by calculating the following empirical adjusted p-value **p_d_** of association (Phipson and Smyth. 2010):

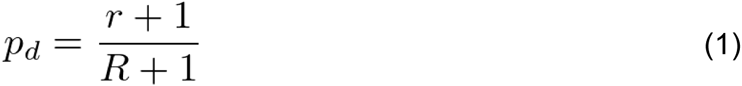

By definition, such an adjusted p-value cannot be smaller than **1/(R+1)**; meaning that a large number of permutations are needed to get good estimates of small adjusted p-values, thereby increasing the computational burden. For instance, reaching p-values of ∼10^-3^ requires thousands of permutations while billions are needed to get p-values of ∼10^-9^. In practice, it is very difficult to go beyond a few thousand permutations genome-wide with this approach, which forces us to work with adjusted p-values in the range of 10^-3^ to 1.0. To alleviate this limitation, we improved the direct permutation scheme with two complementary methods (Methods Sections 4-5).

### 4. Adaptive permutation scheme

From equation (1), one can see that good estimation of insignificant adjusted p-values can be achieved with few permutations while many more are needed to estimate highly significant ones. Therefore, we implemented an alternative permutation scheme that adapts the number of permutations to the significance level of the variant-phenotype pairs (Hubner et al. 2005). The resulting approach saves time at insignificant hits and invests more for significant ones, thereby maintaining a reasonable overall computational cost. Specifically, this adaptive scheme permutes **P** until a given number **B** (typically 100) of null correlations stronger than the observed one are found. To prevent this strategy running too long for the most significant variant-phenotype pairs, the algorithm cannot perform more than **M** (typically 100,000) permutations in total. Then, an adjusted p-value of association for a candidate QTL is derived using:

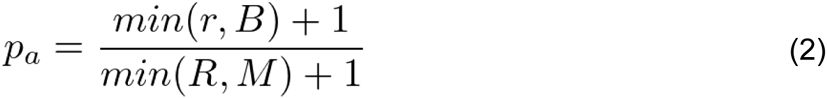

This strategy still remains unable to provide adjusted p-values below **1/(M+1)**, though **M** can scale up to 100,000 in practice and thus provide good estimations for adjusted p-values down to ∼10^-5^.

### 5. Beta approximation

To provide adjusted p-values at any significance level without actually performing all required permutations, we developed an approximation method based on the beta distribution. It is well established that order statistics of independent uniformly distributed random variables are beta-distributed (Jones, 2009). Therefore, we hypothesized that the p-values obtained through permutations are also beta-distributed (Dudbridge & Koeleman, 2004). More formally, the **k^th^** smallest value obtained when independently drawing **n** times from the uniform is distributed as:

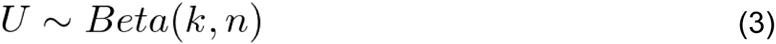

In our particular problem, we propose to model the smallest nominal p-value coming from **L** tests performed in a permutation pass as a beta distributed random variable with shape parameters **k=1** and **n=L**. However, given that nearby variant sites usually exhibit some relatively high degree of correlation (LD), the **L** tests performed are not independent, implying that the effective number of tests **n** is lower than the actual number **L** of variants in cis. Instead of fixing the **k** and **n** parameters a priori, we use a more flexible approach in which the parameters are estimated by maximum likelihood (Galwey. 2009). Specifically, we perform **R** permutations to generate a null set of p-values **{p_1_, …, p_R_}** and then estimate **k** and **n** by maximizing the following log-likelihood:

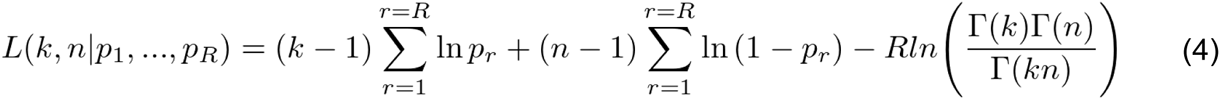

Note that this maximization is done using standard numerical methods implemented in GNU Scientific Library (GSL). The underlying idea of this approach is to characterize the extreme tail of the null distribution without directly sampling from it, something that would entail a huge computational burden. Finally, we can approximate an adjusted p-value **p_b_** from the best nominal p-value **p_n_** and from the maximum likelihood (ML) fitted beta distribution with:

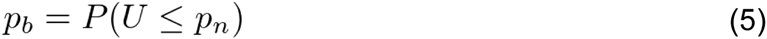

### 6. False discovery rate

Since thousands of molecular phenotypes are tested genome-wide, a false discovery rate correction is commonly applied. This estimates the proportion of false positive findings, known as the FDR, by comparing the number of hits declared to the number that would be expected by chance. The Benjamini-Hochberg (BH) procedure (Benjamini and Hochberg. 1995) is one way of controlling the number of false positive results. However, this is too conservative in most of the QTL studies where we expect a substantial fraction of the phenotypes to be affected by genetic variants. To account for this, it is recommended instead the use of the FDR procedure described by Storey and Tibshirani (ST) (Storey and Tibshirani. 2003) which fits particularly well in this context. The ST procedure assumes that the set of association tests originates from a mixture of both the null and the alternative hypothesis and estimates **π_0_**, defined as the proportion of hypotheses for which the null is true. Implicitly, the BH procedure assumes **π_0_** is 1 whereas the ST procedure learns it from the data, resulting in more statistically significant hits. Of note, the adjusted p-values provided by FastQTL allow the users to easily apply whichever multiple testing correction they favour, from FDR to Bonferroni, since it provides adjusted p-values well calibrated on the full p-value range.

### 7. Implementation

FastQTL implements in C++ Methods Sections 1-5 to provide an adjusted p-value per phenotype. A FDR procedure (Methods Section 6) is then straightforward to apply on the FastQTL output. In addition to the functionalities described above, FastQTL also implements some additional useful features worth mentioning here:

1. To make the method more robust to outliers in the phenotype data, FastQTL has an option that allows to quantile normalize the phenotype quantifications prior to any analysis. This ensures that phenotype quantifications are normally distributed with mean 0 and standard deviation 1. Quantile normalization is implemented as in the R/rntransform function of the GenABEL package (Aulchenko et al., 2007).
2. Confounding factors such as population stratification and experimental batch effects have to be considered to prevent spurious associations. To do so, FastQTL can residualize both the genotypes and the phenotypes for quantitative and/or qualitative covariates prior to any association testing.
3. FastQTL uses standard file formats: genotype dosages and phenotype quantifications are specified in Variant Calling Format (http://vcftools.sourceforge.net/specs.html) and UCSC BED format (http://genome.ucsc.edu/FAQ/FAQformat.html#format1), respectively. All files are required to be indexed with Tabix (Li., 2011) to enable fast retrieval of specific genomic regions.
4. To split a genome-wide analysis into non-overlapping chunks and to run each on a distinct CPU core, FastQTL includes a set of user-friendly options. It can either split the data into a given number of chunk (**--chunk 12 200** to run chunk 12 out of 200) or focus on a particular user defined genomic region (**--region 20:1-1000000**). The phenotype and genotype data included in the genomic region is then automatically extracted from the cis-window size and analyzed. A simple loop going through all possible chunks allows the user to submit the full analysis on a compute cluster or server. FastQTL example command lines to perform a genome-wide analysis or a deep interrogation of a specific region are shown in **command 1 and 2**, respectively.

## Results

To perform a comprehensive evaluation of FastQTL, we used RNA-seq and genotype data produced by both the Geuvadis (Lappalainen et al., 2013; **supplementary material 1**) and the GTEx consortia (GTEx Consortium, 2015; **supplementary material 2**), two of the largest eQTL studies. This comprises a total of 10 distinct data sets with between 14K to 35K quantified genes and 6.8M to 10.8M variant sites for 83 to 373 samples (**supplementary table 1**).

In the context of this work, the two parameters of the beta distribution, **k** and **n**, can be interpreted as the rank of the associated variant and the effective number of independent tests performed in cis, respectively. We looked at the ML estimate distributions of these parameters across all genes in the GEUV_EUR data set and find first that parameter **k** values tend to center around 1.0, in line with what is expected for the top variant (figure 1a). Second, we find that the parameter **n** values show high dispersion (figure 1b) and are consistently smaller than the actual number of variants being tested in cis (figure 1c); both suggesting that the beta distribution captures well the redundancies between variants, a consequence of LD. This also highlights the importance of performing permutations instead of using a Bonferroni correction based on the number of variants, which would result in a substantial proportion of false negative results.

**Figure 1.**
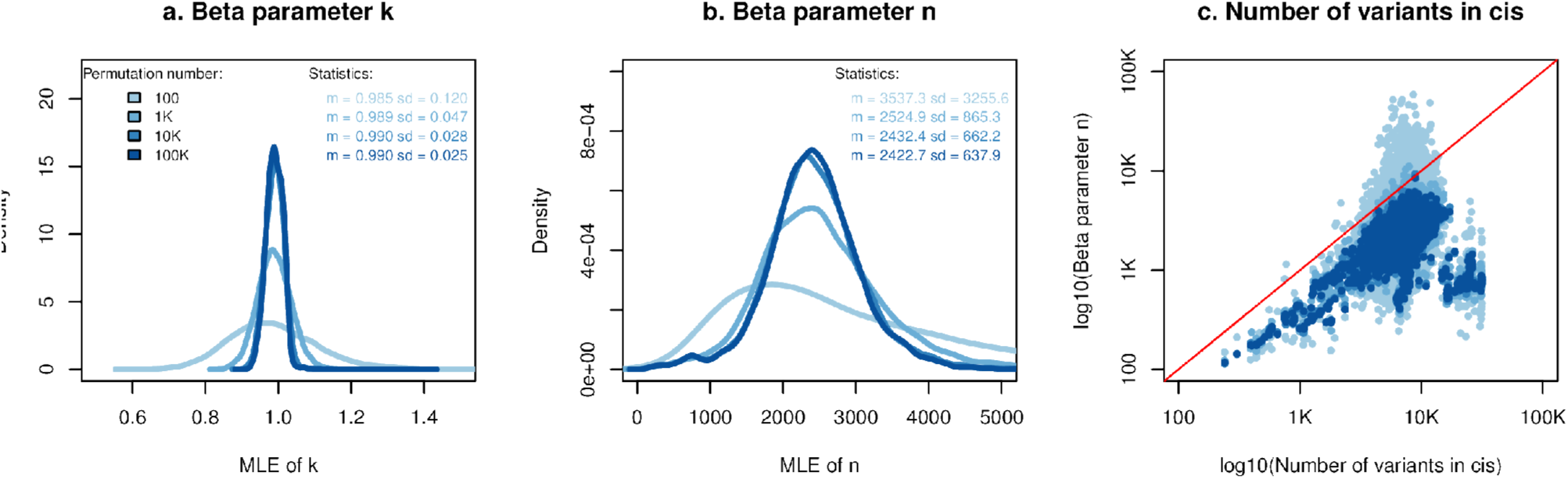
<u>Beta distribution parameters values</u>. Panels (a) and (b) show density plots of the **k** and **n** parameter ML estimates made from 100, 1K, 10K and 100K permutations on GEUV_EUR. Panel (c) shows a scatter plot of the number of variant sites tested per gene (cis-window +/- 1Mb of the TSS) against the **n** parameter ML estimates made from 100, 1K, 10K and 100K permutations on GEUV_EUR (with same color code than for panels (a) and (b)).

Then, we checked whether the null p-values coming from permutations are beta distributed again in the GEUV_EUR data set. To do so, we (1) stored for each phenotype the best p-values obtained from 1,000 permutations as observations, (2) estimated **k** and **n** by ML from the 1,000 resulting p-values, (3) simulated 1,000 p-values from the newly parameterized beta distribution as expectations, and (4) compared both observations and expectations to assess their goodness-of-fit visually (QQ-plots) and statistically (one sample Kolmogorov-Smirnov test). Overall, we find very high degrees of concordance between results; both when pooling all genes together (figure 2a) and also when looking at each gene individually (figure 2b). We only find that the Beta distribution is not a good fit for 2 genes out of the 13,703 tested (figure 2c), these discrepancies are likely due to the stochastic nature of the simulations made in step (3). All this shows that the beta distribution is a good fit for the smallest null p-values generated via permutations and therefore a good candidate to model the permutation process outcome.

**Figure 2.**
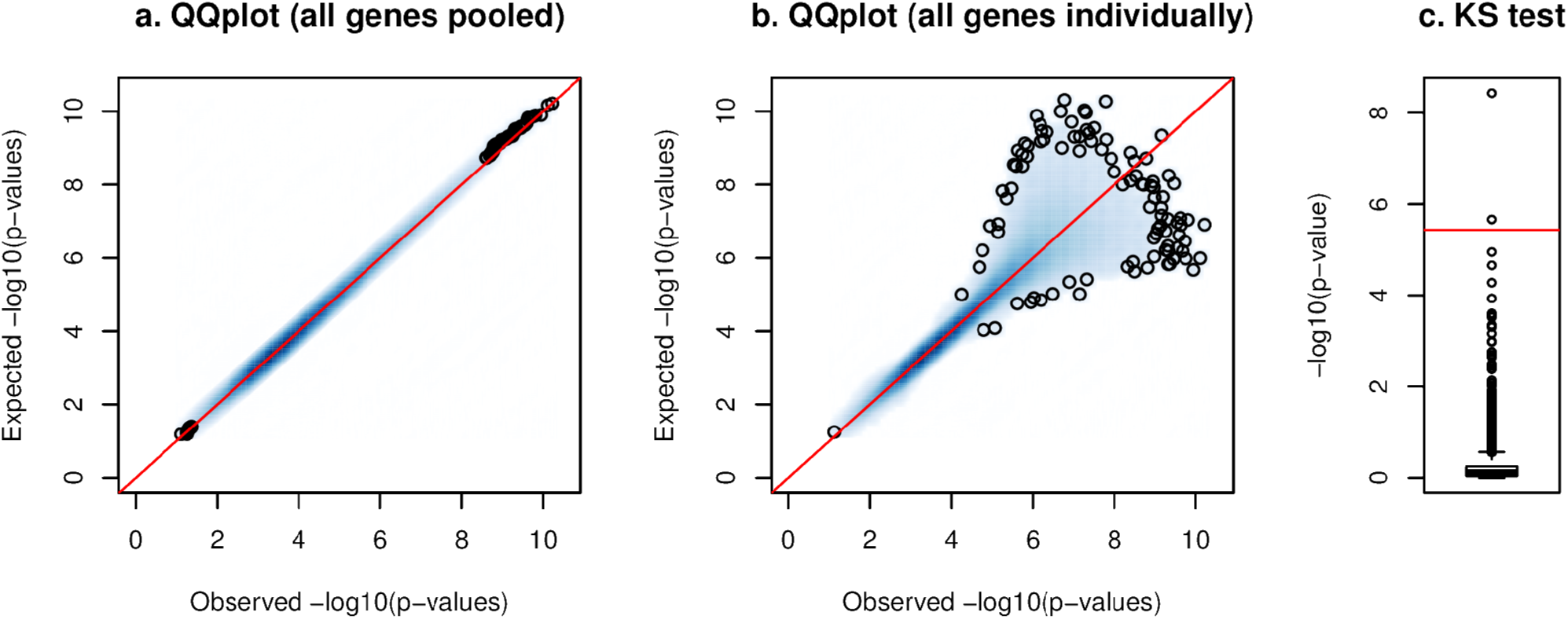
<u>Null p-values are beta-distributed.</u> Panels (a) and (b) show Quantile-Quantile plots of the best p-values obtained through 1,000 permutations (observed) of the GEUV_EUR data set against simulated p-values sampled from the fitted beta distributions (expected). Expected p-values are plotted against the observed ones for all genes pooled together in panel (a) and for each gene separately in panel (b). Panel (c) shows the Kolmogorov-Smirnov (KS) test -log10 p-values comparing observations and expectations for each gene. The red line shows the Bonferroni significance threshold for 13,703 genes tested.

We next checked that the adjusted p-values produced via beta approximation are well calibrated by comparing them to those directly derived from a large number of permutations. We find a very good concordance on the full p-value range with some deviations within the expected sampling variation range (figure 3a, **supplementary figure 1a-b**). Of note, the beta approximation provides small adjusted p-values that are better calibrated than those provided by the direct method (figure 3b, **supplementary figure 1c-d**) and sometimes not even accessible (i.e. below the lower bound implied by the number of permutations); the smallest adjusted p-value estimated using the GEUV_EUR dataset is in the order of ∼10^-128^ (**supplementary figure 2**). Therefore, we subsequently estimated the number of permutations that the direct method needs to reach the same level of calibration as the beta approximation at various significance levels. To do so, we binned the adjusted p-values obtained from beta approximations and estimated for each bin, by exhaustive search, the number of permutations required by the direct method to match the same sampling variation (**supplementary material 3**). We find that this number drastically increases as small-adjusted p-values are targeted (figure 3c). For instance, beta approximations made from 1,000 permutations give adjusted p-values of 10^-4^ as accurately as the direct approach with ∼50K permutations.

**Figure 3.**
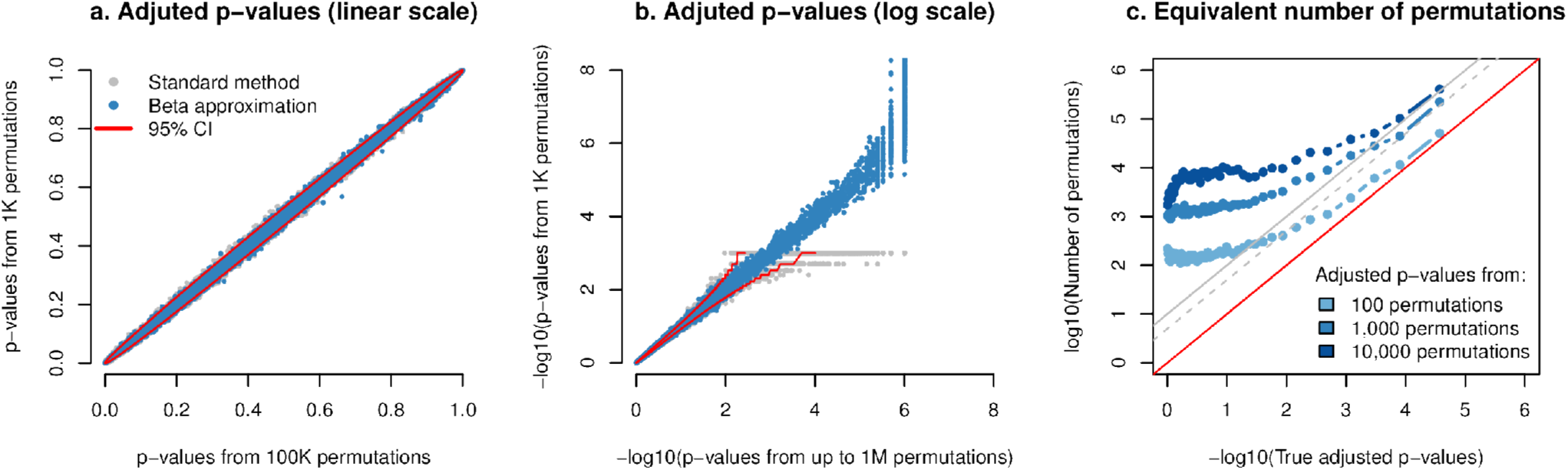
<u>Beta approximated p-values are well calibrated.</u> Panels (a) and (b) show scatter plots of the adjusted p-values obtained from 1,000 permutations via the direct method (in grey) and the beta approximation (in light blue) against those obtained through the standard permutation scheme with 100K permutations (a) or through the adaptive method with up to 1M permutations (c). All this was performed on the GEUV_EUR data set. Adjusted p-values are plotted on both linear (a) and log (b) scales. Expected variation for 1,000 permutations is shown by the 95% confidence intervals in red. Panel (c) shows the equivalent number of permutations required by the direct permutation scheme to reach the same calibration as the beta approximation (from 1,000 permutations) as a function of the adjusted p-value targeted. The dashed and solid gray lines show the expected accuracy of the adaptive permutation scheme that stops when 5 and 10 stronger null signals are found, respectively.

Then, we looked at the downstream impact of the beta approximation on QTL discovery. To do so, we first generated an optimal 5% FDR eQTL set for GEUV_EUR by running 100,000 permutations and then measured the sensitivity/specificity ratios of reasonable FastQTL configurations to recover this optimal set. Specifically, we run from 50 to 5,000 permutations using either the beta approximation or the direct method to compute adjusted p-values. We find that using 1,000 or 5,000 permutations allows us to approximate well the optimal set (figure 4a). We also find that the beta approximation does consistently better than the direct method for the same number of permutations, especially when few permutations are used (50 or 100). As a consequence, using 500 permutations with the beta approximation, for example, has the same accuracy to recover the optimal set as the direct method using 1,000 permutations. Interestingly, the beta approximation using only 50/100 permutations already does very well at recovering the optimal eQTL set.

**Figure 4.**
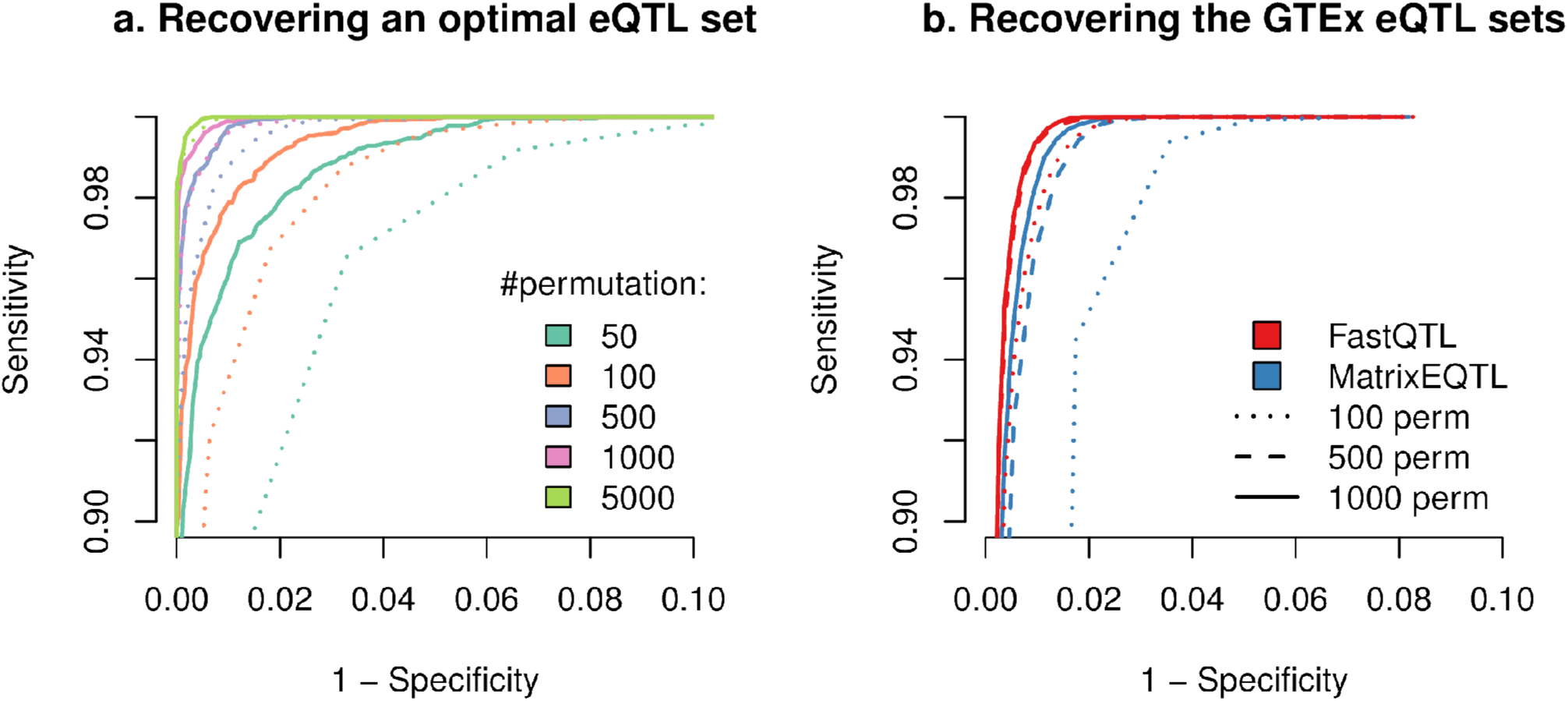
<u>Beta approximation allows fast discovery of eQTLs.</u> Panel (a) shows the sensitivity-specificity ratio to recover an optimal eQTL set derived from 100,000 permutations using reasonable FastQTL runs (beta approximation (solid line) or direct method (dotted line) with 50 to 5,000 permutations). Panel (b) shows the sensitivity-specificity ratio to recover the 9 official eQTL sets released by the GTEx consortium using both Matrix eQTL (direct method) and FastQTL (beta approximation) with 100, 500 and 1,000 permutations.

Finally, we investigated the speed and accuracy with which Matrix eQTL (direct method) and FastQTL (beta approximation) using 100, 500 and 1,000 permutations could reproduce the outcome of the pilot phase of GTEx; a large scale eQTL mapping study. Of note, we run Matrix eQTL in the most highly effective setting we could achieve in order to fully utilize its matrix based design (**supplementary material 4**). Overall, we find that both FastQTL and Matrix eQTL recapitulate the official eQTL set well, especially as the number of permutations increases (figure 4b). For the same number of permutations, we find that the closest eQTL set to the official one is consistently provided by FastQTL. Again, it also performs well even when only 100 permutations are used to fit the beta distributions. To process all 9 data sets with 1,000 permutations, FastQTL requires ∼191 CPU hours which is ∼16 times faster than running the same number of permutations with Matrix eQTL (table 1). When using only 100 permutations, this is reduced to only ∼33 CPU hours.

**Table 1.** <u>FastQTL is fast.</u> This table shows the running times in CPU hours to produce the results shown in figure 4b; 9 GTEx data sets (column 1) processed with 1,000 Matrix eQTL permutations (column 2) and FastQTL with 1,000 (column 3), 500 (column 4) and 100 permutations (column 5). Total running times for all 9 data sets together are shown in the last row.

|  | Matrix eQTL | FastQTL |  |  |
| --- | --- | --- | --- | --- |
| #permutations | 1000 | 1000 | 500 | 100 |
| GTE<sub>x</sub>_AS | 337.4 | 19.0 | 10.8 | 3.5 |
| GTE<sub>x</sub>_AT | 330.3 | 21.0 | 10.8 | 3.6 |
| GTE<sub>x</sub>_HLV | 312.4 | 15.0 | 8.0 | 3.0 |
| GTE<sub>x</sub>_L | 364.9 | 25.6 | 13.1 | 4.0 |
| GTE<sub>x</sub>_MS | 335.9 | 23.6 | 12.6 | 3.9 |
| GTE<sub>x</sub>_NT | 343.8 | 18.4 | 9.5 | 3.4 |
| GTE<sub>x</sub>_SSEL | 349.7 | 20.7 | 10.8 | 3.6 |
| GTE<sub>x</sub>_T | 358.1 | 22.3 | 11.8 | 3.9 |
| GTE<sub>x</sub>_WB | 340.5 | 25.5 | 13.7 | 4.1 |
| ALL | 3073 | 191.1 | 101.1 | 33 |

## Discussion

We present FastQTL, a QTL mapper in cis for molecular phenotypes that implements a new permutation scheme to accurately and rapidly correct for multiple-testing at both the genotype and phenotype levels. FastQTL has several advantages compared to existing methods, making it the ideal candidate to map QTLs for the the coming wave of large scale data sets regrouping many different layers of molecular phenotypes and near complete collection of variant sites. Firstly, permutations are modeled with a beta distribution, parameterized from a relatively small number of permutations. This results in accurate adjusted p-values which could not be feasibly obtained by standard or adaptive permutation analysis; for example down to 10^-128^ in the Geuvadis data set. In practice, having well calibrated adjusted p-values on the full range (0, 1) is of crucial importance to (1) estimate the number of tests made under the null (quantity underlying efficient FDR correction methods) and to (2) meta-analyze multiple QTL studies together. Second, the beta approximation behaves well enough with only 100 permutations to rapidly assess the impact on the analysis of important parameters such as cis-window size and covariates like the number of PEER factors (Stegle et al., 2012). And finally, FastQTL is fast (∼16x faster than Matrix eQTL for the same number of permutations) due to an efficient implementation of linear regressions, optimized C++ code, efficient permutation schemes, and rapid data retrieval from indexed files, while remaining user and cluster friendly. To summarize, FastQTL provides better adjusted p-values than the best method so far, Matrix eQTL with a direct permutation scheme, while being significantly faster. In addition, FastQTL also provides a modular base onto which new functionalities are being implemented, such as fine mapping of causal variants, conditional analysis to discover multiple independent QTLs per phenotype and interaction analysis to discover sex or disease specific QTLs.

## Author contributions

HO, AB & OD designed the method. HO & OD developed the software and carried out the experiments. AAB provided statistical expertise. ETD & OD supervised the research. HO, AAB & OD wrote the paper.

### Funding information

Louis-Jeantet Foundation, NIH-GTEx, European Research Council, Swiss National Science Foundation, SystemsX. Helse S ø r Ø st.

**Command 1**

~~~
(1)  for c in $(seq 1 256); do
(2)         fastQTL      --vcf genotypes.vcf.gz
(3)                      --bed phenotypes.bed.gz
(4)                      --chunk $c 256
(5)                      --permute 1000
(6)                      --output results.$c\.txt.gz
(7)  done
(8)  zcat results.*.txt.gz I gzip -c > results.full.txt.gz
~~~

<u>Genome-wide analysis example.</u> This shows the BASH script needed to run a genome-wide analysis. The genotypes and phenotypes are specified with **--vcf** (line 2) and --**bed** (line 3), respectively. The analysis is split into 256 non-overlapping chunks (lines 1 and 4) and is based on 1,000 permutations (line 5). The full outcome is constructed by concatenating the per-chunk outcomes (line 8).

**Command 2**

~~~
(1)  fastQTL       --vcf genotypes.vcf.gz
(2)                --bed phenotypes.bed.gz
(3)                --region 4:15000000-16000000
(4)                --permute 1000000
(5)                --output results.chr4.15M.16M.txt.gz
~~~

<u>Deep interrogation of a specific region.</u> This shows the BASH script needed to run the analysis of all molecular phenotypes located on chromosome 4 with coordinates between 15Mb and 16Mb (line 3). This analysis relies on 1,000,000 permutations (line 4).

## References

Aulchenko,Y.S. et al. (2007) GenABEL: an R library for genome-wide association analysis. Bioinformatics. 23:1294–6.

Benjamini,Y. and Hochberg,Y. (1995) Controlling the False Discovery Rate: A Practical and Powerful Approach to Multiple Testing. Journal of the Royal Statistical Society. 57:289–300.

Dudbridge,F. and Koeleman,B.P. (2004) Efficient computation of significance levels for multiple associations in large studies of correlated data, including genomewide association studies. Am J Hum Genet. 75:424–35.

Fairfax,B.P. et al. (2012) Genetics of gene expression in primary immune cells identifies cell type-specific master regulators and roles of HLA alleles. Nat Genet. 44:502–10.

Galwey,N.W. (2009) A new measure of the effective number of tests, a practical tool for comparing families of non-independent significance tests. Genet Epidemiol. 33:559–68.

GTEx Consortium. (2015) The Genotype-Tissue Expression (GTEx) pilot analysis: multitissue gene regulation in humans. Science. 348:648–60.

Hubner,N. et al. (2005). Integrated transcriptional profiling and linkage analysis for identification of genes underlying disease. Nat Genet. 37:243–53.

Jones,M.C. (2009) Kumaraswamy’s distribution: a beta-type distribution with some tractability advantages. Statistical Methodology. 6:70–81.

Lappalainen,T. et al. (2013) Transcriptome and genome sequencing uncovers functional variation in humans. Nature. 501:506–11.

Li,H. (2011) Tabix: fast retrieval of sequence features from generic TAB-delimited files. Bioinformatics. 27:718–9.

Maurano,M.T. et al. (2012) Systematic localization of common disease-associated variation in regulatory DNA. Science. 337:1190–1195

Montgomery,S.B. et al. (2010) Transcriptome genetics using second generation sequencing in a Caucasian population. Nature. 464:773–7.

Nica,A.C. et al. (2010) Candidate causal regulatory effects by integration of expression QTLs with complex trait genetic associations. PLoS Genet. 6(4):e1000895.

Ongen,H. et al. (2014) Putative cis-regulatory drivers in colorectal cancer. Nature. 512:87–90.

Phipson,B. and Smyth,G.K. (2010) Permutation P-values should never be zero: calculating exact P-values when permutations are randomly drawn. Stat Appl Genet Mol Biol. doi: 10.2202/1544-6115.1585.

Shabalin,A.A. (2012) Matrix eQTL: ultra fast eQTL analysis via large matrix operations. Bioinformatics. 28:1353–8.

Stegle,O. et al. (2012) Using probabilistic estimation of expression residuals (PEER) to obtain increased power and interpretability of gene expression analyses. Nat Protoc. 7:500–7.

Storey,J.D. and Tibshirani,R. (2003) Statistical significance for genomewide studies. Proc Natl Acad Sci. 100:9440–5.

